# Sensitive detection of highly fragmented cytomegalovirus nucleic acid in human cfDNA

**DOI:** 10.1101/748434

**Authors:** Vikas Peddu, Benjamin T. Bradley, R. Swati Shree, Brice G. Colbert, Hong Xie, Tracy K. Santo, Meei-Li Huang, Edith Y. Cheng, Eric Konnick, Stephen J. Salipante, Keith R. Jerome, Christina M. Lockwood, Alexander L. Greninger

## Abstract

Congenital human cytomegalovirus (CMV) infections are the leading cause of newborn hearing and central nervous system impairments worldwide. Currently, routine prenatal screening for congenital CMV is not performed in the United States and confirmation in suspected perinatal cases requires invasive sampling by amniocentesis. We hypothesized that detection of CMV from maternal cell-free DNA (cfDNA) plasma could provide a non-invasive indicator of congenital CMV infection. We analyzed sequence data from 2,208 individuals undergoing routine non-invasive prenatal aneuploidy screening at the University of Washington. CMV reads were identified in 117 (5.3%) samples. Positive samples were stratified based on CMV reads per million sample reads (RPM), resulting in ten samples being classified as strong positive (RPM > 0.3) and 107 as intermediate positive (0.01<RPM<0.3). Subsequent qPCR testing identified CMV in 9/10 strong positive samples and 2/32 intermediate positive samples. Median cfDNA insert size derived from CMV was significantly shorter than cfDNA derived from human chromosomes (103 vs 172 bp, p<0.0001), corresponding to the 3^rd^ percentile of human cfDNA insert size. In addition, CMV cfDNA fragment lengths were distributed over a wider range than human cfDNA reads. These studies reveal the highly fragmented nature of CMV cfDNA and offer precise measurements of its length: these features likely explain discrepancies in serum CMV viral loads measurements determined by different qPCR assays, despite widespread efforts to standardize results. More work is required to determine how detection of CMV from maternal cfDNA can be best used as tool for congenital CMV screening or diagnosis.

## Introduction

Human cytomegalovirus (CMV) is the most common cause of congenital defects in the United States and affects roughly 0.5-1.3% of live births worldwide (1, 2), causing more congenital disease than all disorders tested for in newborn screening combined (4). Congenital CMV is associated with a variety of late-onset permanent disabilities such as hearing loss, microcephaly, vision defects, and intellectual deficits (3). Primary maternal CMV infection is associated with a 40% risk of fetal infection, whereas recurrent infection has an approximately 1% transmission rate (3). The high prevalence of recurrent infection means that recurrent disease is the cause of nearly three-quarters of congenital CMV cases (3). Despite the incredible burden of congenital disease, existing methods to screen for fetal CMV infection are limiting in multiple important respects, including a lack of sensitivity and delayed identification of affected patients.

Due to the significant risks posed by primary congenital CMV, prenatal CMV screening via maternal serology has been proposed as an attractive approach to limit the incidence of disease in this population (5). Yet, a variety issues can arise during interpretation of serology results, making diagnosis challenging (6–9). Additionally, the utility of serologic screening is limited to mothers with primary infections, thus excluding the substantial majority of congenital CMV cases that are associated with maternal reactivation or reinfection. Consequently, the current gold standard for diagnosing congenital CMV infection prenatally is amniocentesis, an invasive procedure best performed after 21 weeks gestation. Because of the risks associated with this procedure, amniocentesis is considered following radiologic and/or serologic evidence for congenital disease (10–12).

If not diagnosed antenatally, additional cases of congenital CMV are identified by testing for viral DNA in newborn saliva or urine within the first three weeks following a failed hearing screen or other concerning clinical characteristics (13). Although this is an important strategy to enable early intervention, the underlying developmental abnormalities are already established at the time of testing and only demonstrate a modest response to antiviral treatment (14). Further limiting the impact of this approach is the observation that over half of congenital CMV cases manifest symptoms months to years after birth (15). In cases where delayed symptom onset is suspected, the diagnosis of congenital CMV infection is limited to retrospective testing of stored heel stick blood spots since PCR testing for CMV viremia after three weeks of life cannot exclude disease secondary to postnatal infection (16). In summary, the current landscape of congenital CMV screening is hindered by insensitive tools and delayed identification of cases.

Non-invasive prenatal testing (NIPT) via maternal cell-free DNA (cfDNA) has already revolutionized the ability to screen for fetal aneuploidies, subchromosomal copy number alterations, and other genetic diseases (17, 18). CMV have previously been detected in cfDNA sequencing data, including NIPT (19). At our institution, we have performed clinical screening for fetal aneuploidies by cfDNA since May 2017. As currently available non-invasive screening methods for congenital CMV infection are fraught with error and uncertainty, leveraging prenatal cfDNA sequencing would provide a new non-invasive approach for examining CMV disease burden and predicting clinical outcomes.

## Methods

### Study population

We included all maternal plasma samples collected between May 2017 to November 2018 for clinically-indicated aneuploidy screening performed at the University of Washington Department of Laboratory Medicine. The 2,208 cfDNA samples in our cohort were derived from pregnant women in the University of Washington (UW) Medicine network. Of these, 727 tests were performed during validation and 1,481 tests were performed during the clinical implementation phase. Metadata was available for the 1,325 patients (1,481 tests) screened during the clinical implementation phase (Supplementary Table 1). A minimum gestational age of 10 weeks was required for testing. Following University of Washington Institutional Review Board review and approval, maternal and neonatal clinical histories were gathered for those samples with a CMV cfDNA reads per million sample reads (RPM) > 0.3 (10 samples, 9 patients) or those with a CMV cfDNA RPM < 0.3 and a positive qPCR result (2 samples, 2 patients). We collected maternal age, gravidity, parity, preexisting comorbidities, and results from ultrasound studies. Neonatal information included gestational age at birth, birth weight, APGAR scores, and mode of delivery. Additional information gathered from the antenatal period included admission to the neonatal intensive care unit (NICU), length of stay, and any infectious disease testing performed. Given the available cfDNA data, we also included maternal and placental cfDNA fractions and the number of total reads.

### Non-invasive cell-free DNA sequencing

cfDNA reads from maternal plasma were generated through a validated, laboratory-developed method used to screen for fetal aneuploidies and copy number alterations. For sample preparation, whole blood from Streck (BCT1) tubes was centrifuged and plasma was isolated as per the package insert. cfDNA was extracted from plasma using the QIAsymphony Circulating DNA Kit. Following measurement of the DNA concentration in the eluate, next-generation sequencing library preparation was performed on the BioMek 4000 using the KAPA HyperPrep kit for adapter and index ligation. The library was purified using the Agencourt AMPureXP kit prior to amplification. Following amplification, the library was purified on the Agilent BRAVO workstation using AMPure beads. Sample pools were created using an equimolar strategy and diluted to 1nM. Sequencing was performed using an Illumina NextSeq 500 using the High Output 75 cycle kit, with a 37bp paired-end read configuration.

### Cell-free DNA bioinformatics pipeline for detection of CMV

Paired-end 37bp cfDNA reads were aligned against the human herpesvirus 5 Merlin strain reference genome (NC_006273.2) using bowtie2 (flag: --local –no-unal). The resulting alignment file was filtered to exclude any aligned reads with fewer than 34 exact matches to the reference. Reads aligning to CMV by bowtie2 were then also confirmed via BLASTn alignment to the reference genome with a minimum evalue of 1e-5. CMV levels by cfDNA sequencing were quantified as CMV-specific RPM. A threshold of greater than or equal to 0.3 RPM was set as strong positive while any value greater than zero and less than 0.3 RPM was set as intermediate positive. Fragment length was calculated from the insert size column of the paired-end bam file for sample 121R04 after removal of duplicates using Picard’s MarkDuplicates command (http://broadinstitute.github.io/picard). Statistics and graphical plotting were performed in R using ggplot2, ecdf, t.test, and Kolmogorov–Smirnov statistical tests (20). For median cfDNA insert size comparison and cumulative frequency distribution graphing, human reads were randomly downsampled to the same number as the CMV reads and statistical tests performed over 10,000 iterations.

### Cell-free DNA bioinformatics pipeline for detection of inherited chromosomally integrated HHV-6 (iciHHV-6)

All cfDNA sequences were aligned against telomere-trimmed versions of the HHV-6A (NC_001664.4) and HHV-6B (AF157706.1) reference genomes using the same bowtie2 options as above. Any samples detected with non-repeat reads aligning in the HHV-6A or HHV-6B U38- U100 region were selected for further analyses.

These positive cfDNA files were then also aligned to portions of the human genes EDAR (NM_022336.4) and beta-globin (AH001475.2) that were trimmed of human repeats (Supplemental Figure 1) using the same bowtie2 options specified above. Normalized depth of coverage was calculated by dividing all values by the highest RPKM determined and multiplying by 100 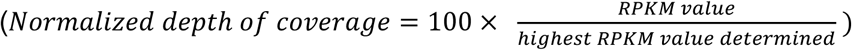. RPKM values were calculated for each sample as 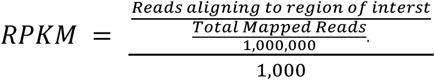.

### PCR detection of CMV from cell-free DNA

The 10 strong positive samples, a random selection of 32 intermediate positives, and 25 run-matched negative controls were analyzed by qPCR to confirm CMV detection. Copy number for positive samples are presented in Supplementary Table 2. Remnant cfDNA from the original maternal plasma extraction was diluted to obtain 50µL of DNA. 15µl of sample was loaded into a 96 well plate with 17.5 µl of Bio-Rad Ssoadvanced Universal Probes Supermix. Plates were sealed, mixed, and vortexed prior to amplification on an Applied Biosystems QuantStudio 7 Flex for 46 cycles at 50C for 2 minutes, 95C for 23 seconds, and 60C for 30 seconds. Samples were run using primers and probes specific for the gB and IE EX-4 regions of CMV and human β –Globin (Supplementary Table 3). Copy number per milliliter was calculated using a standard curve. Cycle thresholds were compared to the RPM values generated from the bioinformatics pipeline using linear regression.

## Results

### Characteristics of study population and testing performance

The median gestational age for the 1,322 patients (1,481 samples) tested during the clinical implementation phase was 13 weeks and 6 days. Average maternal age was 34 years 6 months. The most common indication for testing was advanced maternal age, accounting for 58.3% of cases. Over the study period examined, a total of 35 aneuploidies were detected. For specimens with CMV detected by cfDNA and qPCR, maternal and neonatal outcomes were obtained via medical record review when available. Demographic features of patients tested are listed in Supplemental Table 1 and a comparison between the CMV cfDNA positive and negative populations are presented in Supplemental Figure 1.

For the 2,208 samples in this study, the median number of total reads per sample was 26,563,081 (central 95^th^ percentile of 6,770,026 to 70,439,008). Variation in read depth was related to the number of samples batched per run, ranging from five to twenty-seven. A fetal fraction greater than 4% is required for clinical reporting of fetal aneuploidies; however, for this study, all results were analyzed regardless of fetal fraction.

### Cytomegalovirus detection in cfDNA

A total of 117/2208 samples (5.3%) contained at least one read mapping to CMV. 107 were subsequently defined as intermediately positive (0<RPM<0.3) and 10 were defined as strongly positive (RPM>0.3). The RPM values of the CMV-positive cfDNA samples are shown in Figure 1. Verification by qPCR testing was performed on all 10 strongly positive samples, a subset of intermediately positive (n=32), and run-matched negative controls (n=25). qPCR detected copies of CMV in 9/10 high positive samples and 2/32 intermediate positives. The control samples were appropriately negative for CMV and positive for beta-globin. We next compared the calculated CMV cfDNA RPM to viral load by qPCR. Linear regression analysis demonstrated a weak positive correlation between cfDNA CMV reads and viral load by qPCR (R^2^= 0.42) over the concentrations examined (Figure 2).

**Figure 1.**
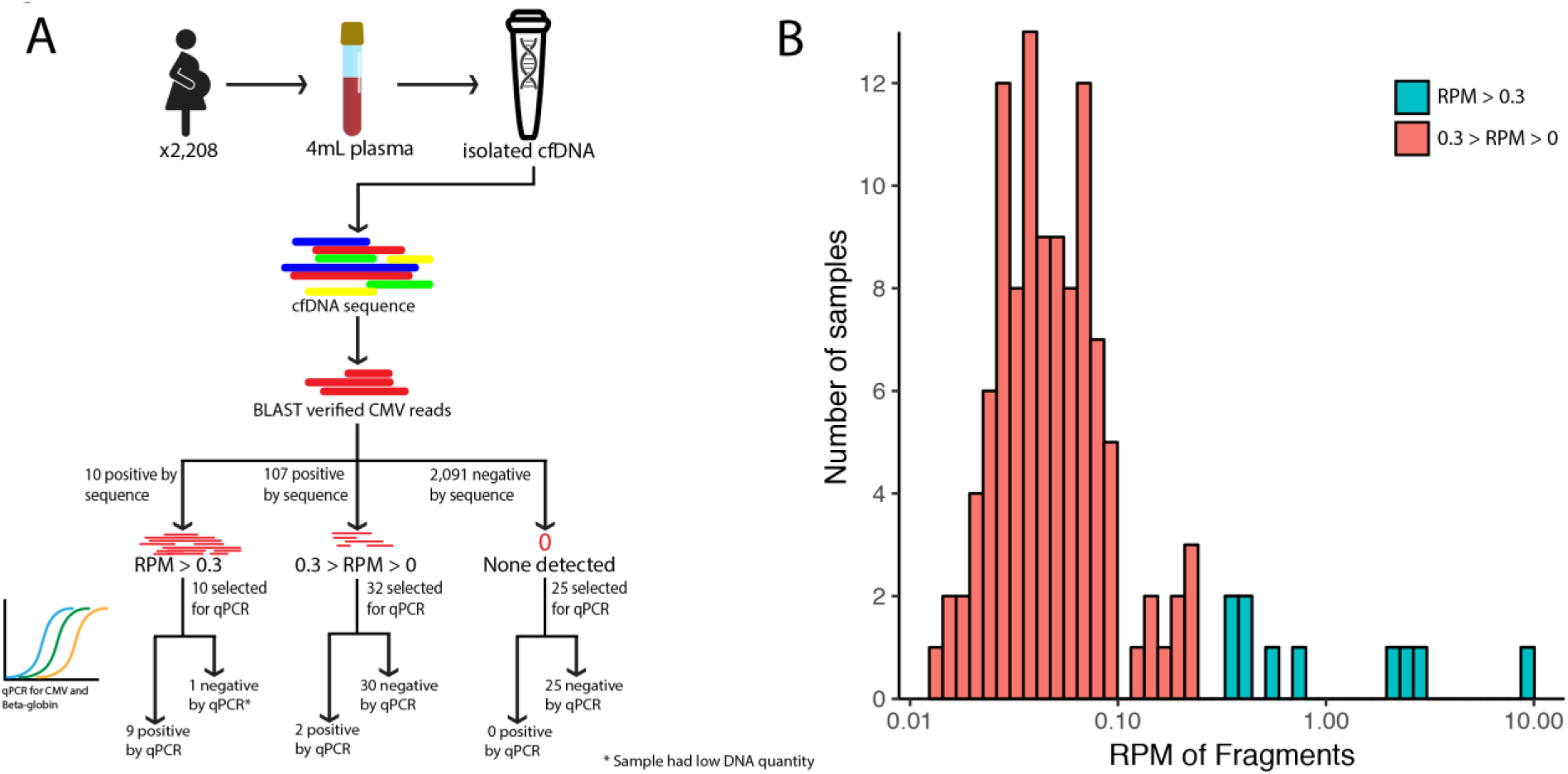
cfDNA pipeline and results. A) Specimen handling for detecting CMV cfDNA. Following library sequencing, data was aligned to the human herpesvirus 5 Merlin strain reference genome (NC_006273.2). A subset of samples was tested via qPCR. B) CMV read distribution by sample. An arbitrary threshold of 0.3 CMV reads per one million reads (RPM) was set to classify specimens as strong (blue) or intermediate positive (red).

**Figure 2.**
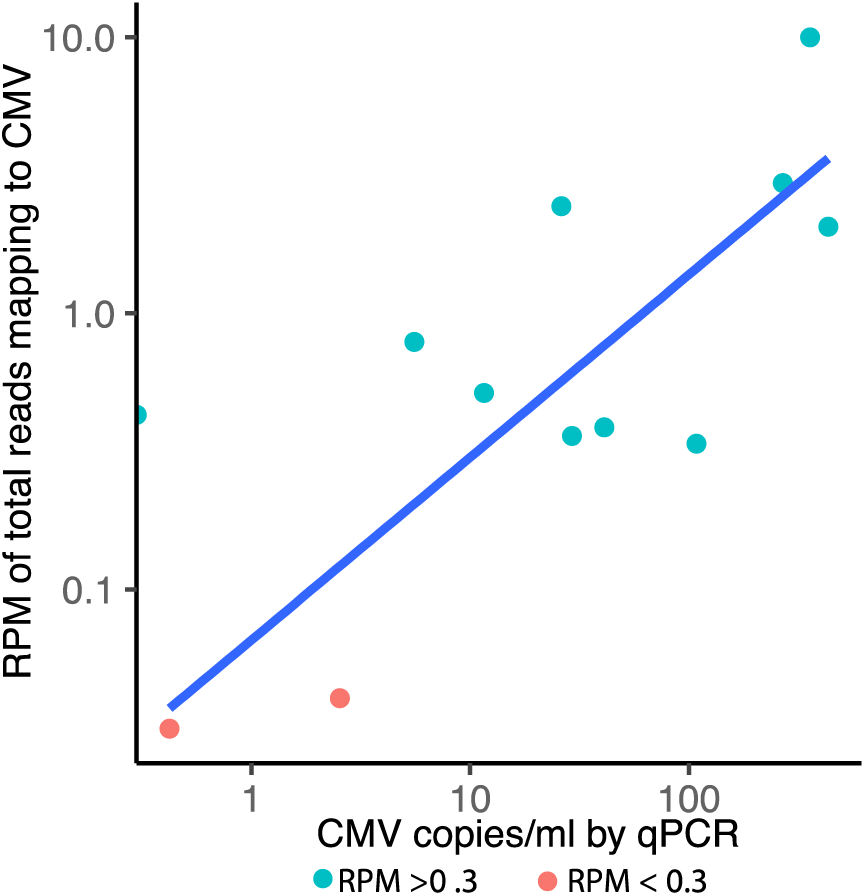
CMV cfDNA reads per million (RPM) correlate with DNAemia calculated by qPCR. For samples identified with positive CMV cfDNA reads and positive qPCR, comparing the values via scatter plot demonstrates a positive correlation between results (R^2^=0.46).

### Short fragment length of CMV cfDNA

In order to attain sufficient reads to estimate the fragment length of CMV cfDNA, we took the cfDNA sample (121R04) with the highest percent CMV measured by sequencing and sequenced it to a total depth of 498 million paired-end reads. Of these, 4,098 paired reads mapped to CMV by bowtie and BLASTn analysis, of which 2,055 fragments remained after deduplication. Reads obtained from our pipeline generally aligned across the length of the CMV genome but had a noticeable lack of coverage in the RL12-RL13-UL1 region (10-13kb locus in the NC_006273.2 reference genome) (Supplemental Figure 2).

We found that the median CMV fragment length in cfDNA was significantly shorter than that of human-derived cfDNA (103 v. 172 bp, p=4.1e-102), placing it at the 3^rd^ percentile of human cfDNA fragment size (Figure 3A). The distribution of CMV cfDNA fragment size [IQR 63-170 bp] was also significantly different than that of human-derived cfDNA [IQR 158-190 bp] (Kolmogorov–Smirnov test, p<2.2e-16) (Figure 3B).

**Figure 3.**
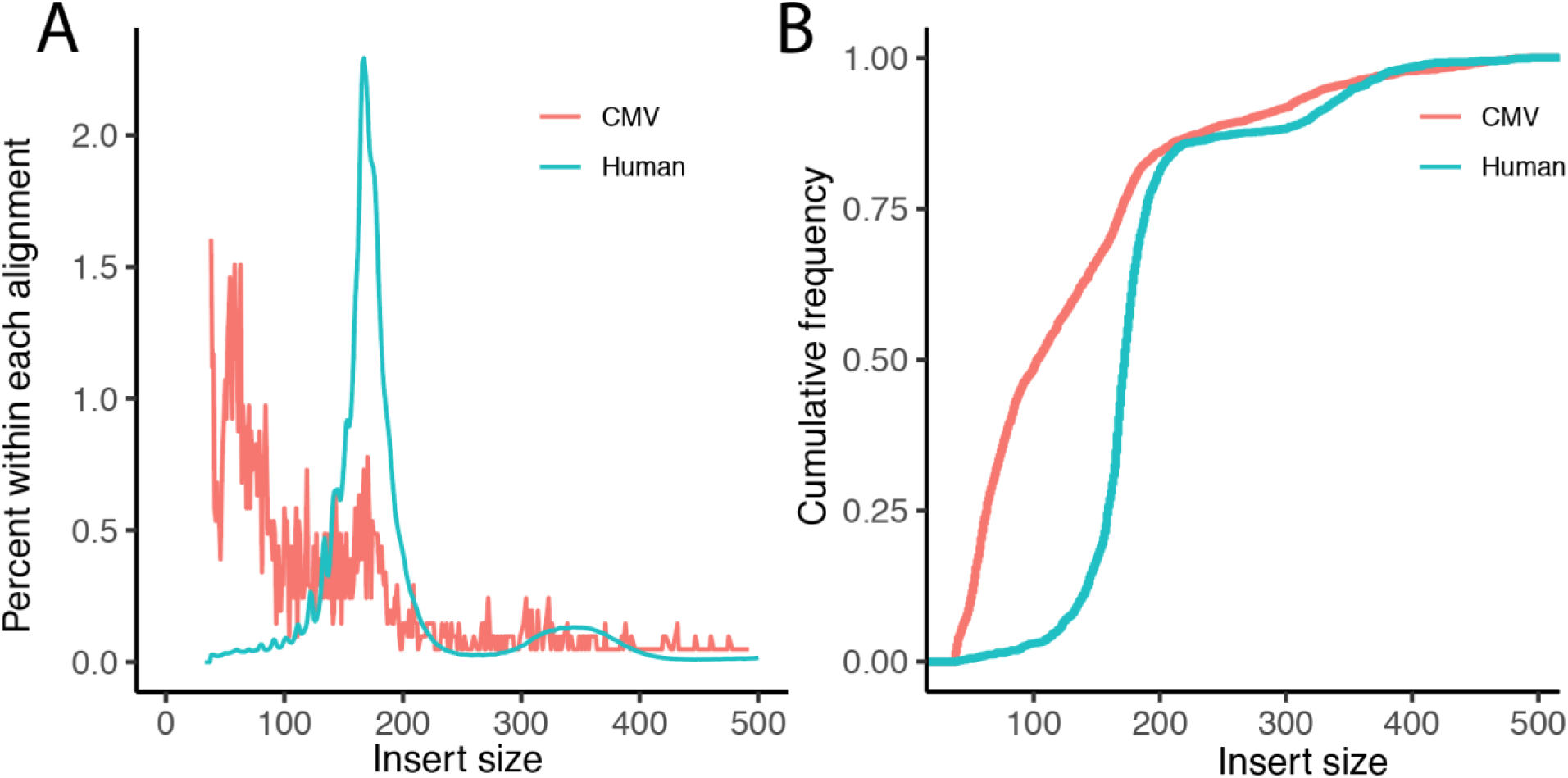
CMV cfDNA insert size from 121R04 is significantly shorter than that of human cfDNA. The median fragment length for CMV cfDNA was 103 [IQR 63-170 bp], while that of human cfDNA was 172 bp [IQR 158-190 bp].

### HHV-6 detection in cfDNA

In order to compare to another betaherpesvirus, we also looked at cfDNA detection of HHV-6. 18 cfDNA samples from 17 different patients had reads aligning to the U38-U100 region of HHV-6A or HHV-6B and were classified as HHV-6 positive. Of these 18 positive samples, 12 had a significantly higher ratio of genomic copies of HHV-6:human genome copies (EDAR or beta-globin), likely consistent with inherited chromosomally-integrated HHV-6 (Figure 4A) (21). The median fragment length of HHV-6 cfDNA by NIPT sequencing across all positive samples was 167 bp [IQR 149-181 bp], approximating that of human-derived cfDNA. When we specifically compared at the fragment length of the twelve high and six low HHV-6 samples compared to human cfDNA, the six low level HHV-6 samples had a shorter fragment length (median 146 bp [IQR 104-176 bp]) than the twelve high level HHV-6 samples (Figure 4B/C). These results are most consistent with a model of normal chromatinization of maternal iciHHV-6 DNA in the high level HHV-6 samples, with the shorter fragments of low level HHV-6 deriving from the placenta due to paternal transmission of iciHHV-6 to the fetus.

**Figure 4.**
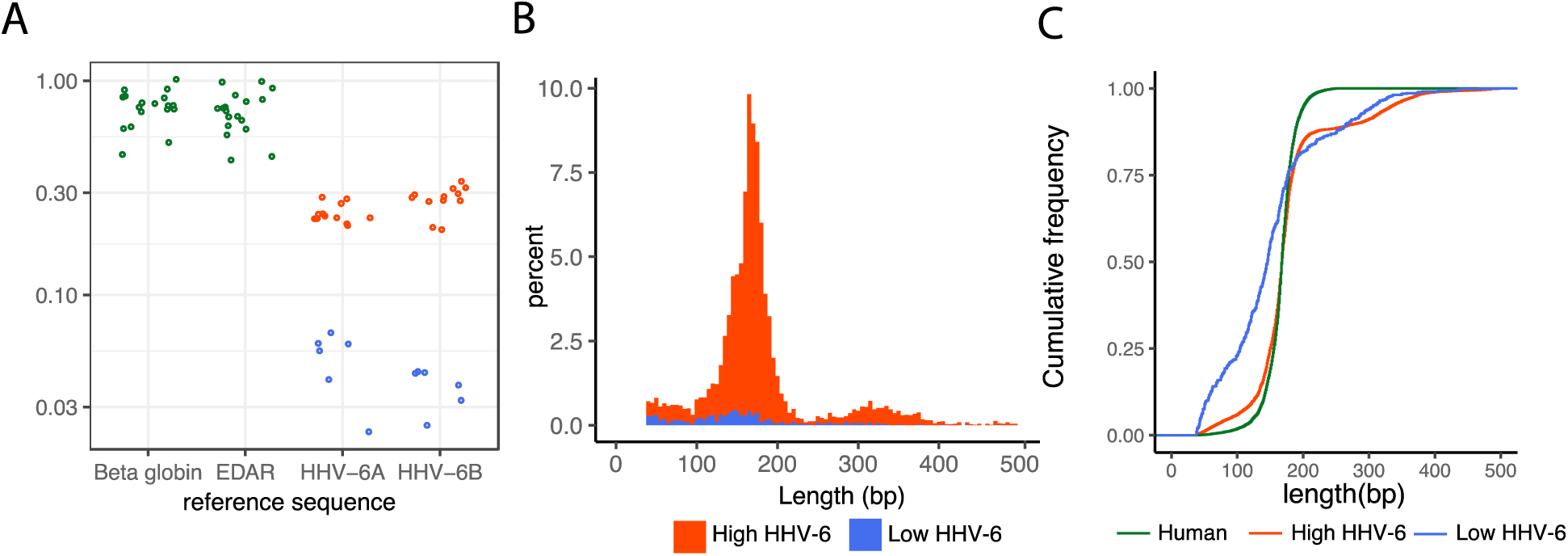
Eighteen samples had HHV-6 cfDNA present, of which twelve had higher levels consistent with iciHHV-6 and six had lower levels relative to human housekeeping genes, beta-globin and EDAR (A). The fragment length distribution of the HHV-6 cfDNA from the high cluster mirrors that of human cfDNA (B). The median fragment length of cfDNA for the high HHV-6 cluster cfDNA was 167 bp [IQR 149-181 bp], while that of low HHV-6 cluster was 146 bp [IQR 104-176 bp] (B and C).

### Clinical outcomes of mother and fetus following CMV cell-free DNA detection

Clinical records for five mothers and five offspring were available from the ten strong positive CMV cfDNA samples (9 patients) and two qPCR-positive, CMV cfDNA intermediate positive samples (2 patients). Clinical records were not available for the remaining cases. Maternal age ranged from 29 to 39 years with a parity range of 1 to 3. Fetal outcomes included elective termination for trisomy 21, preterm delivery at 20 weeks gestation, and three live births. Two of the live births were uncomplicated full-term vaginal deliveries, while the other delivered via Cesarean section at 32 weeks gestation due to decreased fetal movement and non-reassuring fetal status. This preterm infant had low APGAR scores and was admitted to the neonatal intensive care unit for prematurity and respiratory distress. The infant was discharged home after 40 days of hospitalization without further apparent complications. In none of the above cases was CMV PCR performed on the mother or neonate.

## Discussion

Given the global impact and life-long complications of congenital CMV infection, an effective screening strategy to facilitate early intervention is necessary. In exploring the technical feasibility of CMV cfDNA detection, we have discovered that CMV cfDNA exists in circulation at a smaller fragment size than human chromosomally-derived cfDNA. CMV detection from prenatal cfDNA samples sent for aneuploidy testing offers an added benefit over plasma qPCR in that no additional wet lab testing is required.

By establishing the fragment size of CMV cfDNA through ultra-deep sequencing, we have identified a potential mechanism contributing to the variation in CMV levels obtained from different qPCR assays. Despite a recently implemented international standard, variation in plasma CMV DNA levels measured by qPCR across different clinical laboratories has remained high, with variation of up to 2 log_10_ in copy number between assays (24). Amplicon size has been identified as a major contributor to interassay variability, with larger amplicons having a relatively lower IU/ml as compared to smaller amplicons (25). Given that amplicons in CMV qPCR assays range in size from 52 to 340bp, it is reasonable to assume that PCR assays developed for smaller amplicons would more readily amplify CMV cfDNA fragments. Previous studies indicated that CMV DNA in plasma exists as highly fragmented, virion-free DNA, suggesting that the cfDNA fragments measured here may constitute the vast majority of the CMV DNA present in plasma (26, 27). Characterizing CMV cfDNA insert size may also assist in designing target enrichment strategies and improving bioinformatic pipelines for infectious disease screening. The CMV reads obtained from the cfDNA data were generally distributed across the genome without preferential representation of any one region, although curiously low coverage was seen across the RL12-RL13-UL1 gene region, which was also seen in a prior cfDNA study (28). Future studies will be needed to examine more samples to test the generalizability of our single specimen estimation and to explore the effect of cfDNA *in vivo* as it relates to CMV DNA quantitation.

To demonstrate that our approach has the robustness to identify other viral pathogens, we interrogated the cfDNA sequencing data for evidence of iciHHV6. When our cohort of maternal cfDNA samples were analyzed, we identified 18 HHV6-positive samples, 12 of which appeared consistent with maternal iciHHV6 based on fragment length and copy number (Figure 4). Differences in the size distribution of placental and maternal cfDNA may allow for the identification of fetal iciHHV6 in iciHHV6-negative mothers. In the six HHV6-positive samples with an RKPM value below what is expected in iciHHV6, the cfDNA size distribution of the HHV6 reads closely matched that seen in placental cfDNA. We hypothesize the shorter read distribution arising in this population is the result of iciHHV6 from placental DNA fragments.

In our qPCR studies measuring CMV DNAemia, one cfDNA strong positive sample was negative despite a detectable housekeeping control gene. In this sample, the beta-globin signal was 1.0 log_10_ lower than the next lowest positive sample. This finding strongly argues that the negative CMV qPCR result was a function of low input DNA as opposed to a true negative. Consistent with this hypothesis, within the intermediate positive samples only 2 of the 32 tested were positive. The relative insensitivity of qPCR may be related to an insufficient number of DNA fragments in the sample spanning the full length of the amplicon. Alternatively, these false negative qPCR results may be due to the volume of DNA available after cfDNA sequencing. Of note, in our standard qPCR assay we elute 100µl of DNA extracted from 200µL of plasma and perform qPCR on 15µL of the DNA eluate. In the qPCR experiments for this study, our DNA was derived from the eluate of the cell-free extraction method wherein 4mL of whole blood is drawn, extracted, and eluted into 60µl, of which 30µL is used for library preparation. The remaining sample volume available for qPCR from these extractions ranged from 1 to 20µl. We standardized the volume using ultra-pure water and adjusted the calculated DNAemia according to the input volume.

Our study is chiefly limited by the comparatively small number of patients used for CMV cfDNA fragment length estimation and clinical chart review, despite the screening of more than 2,200 specimens. We chose to more deeply sequence one specimen with the highest amount of CMV as recovered by cfDNA sequencing. This may bias our estimation of CMV cfDNA fragment length in the general population, since this specimen had comparatively more reads to CMV as would be predicted by qPCR. Certainly, more cfDNA sequencing in other clinical contexts where CMV detection is critical such as stem-cell or solid organ transplant is required. Because clinical outcomes were only available in five patients, drawing any conclusions in regard to clinical outcomes of a positive CMV cfDNA result is difficult. Maternal data is lacking for six of the strongly CMV cfDNA positive samples due to the mother receiving her subsequent obstetric care at an outside institution. Furthermore, our hospital system is separate from the major pediatric hospital in our area. As a result, we were unable to obtain any significant clinical history or post-natal outcomes for a portion of the strongly positive cases we identified. While at least one case resulted in prolonged hospitalization due largely to prematurity, no CMV testing was performed and imaging studies did not suggest congenital CMV infection. Despite the potential of cfDNA as a diagnostic tool in early detection of congenital CMV, future studies must address whether a level of CMV cfDNA predictive of congenital infection exists.

## Supplemental Tables

**Supplemental Table 1.**
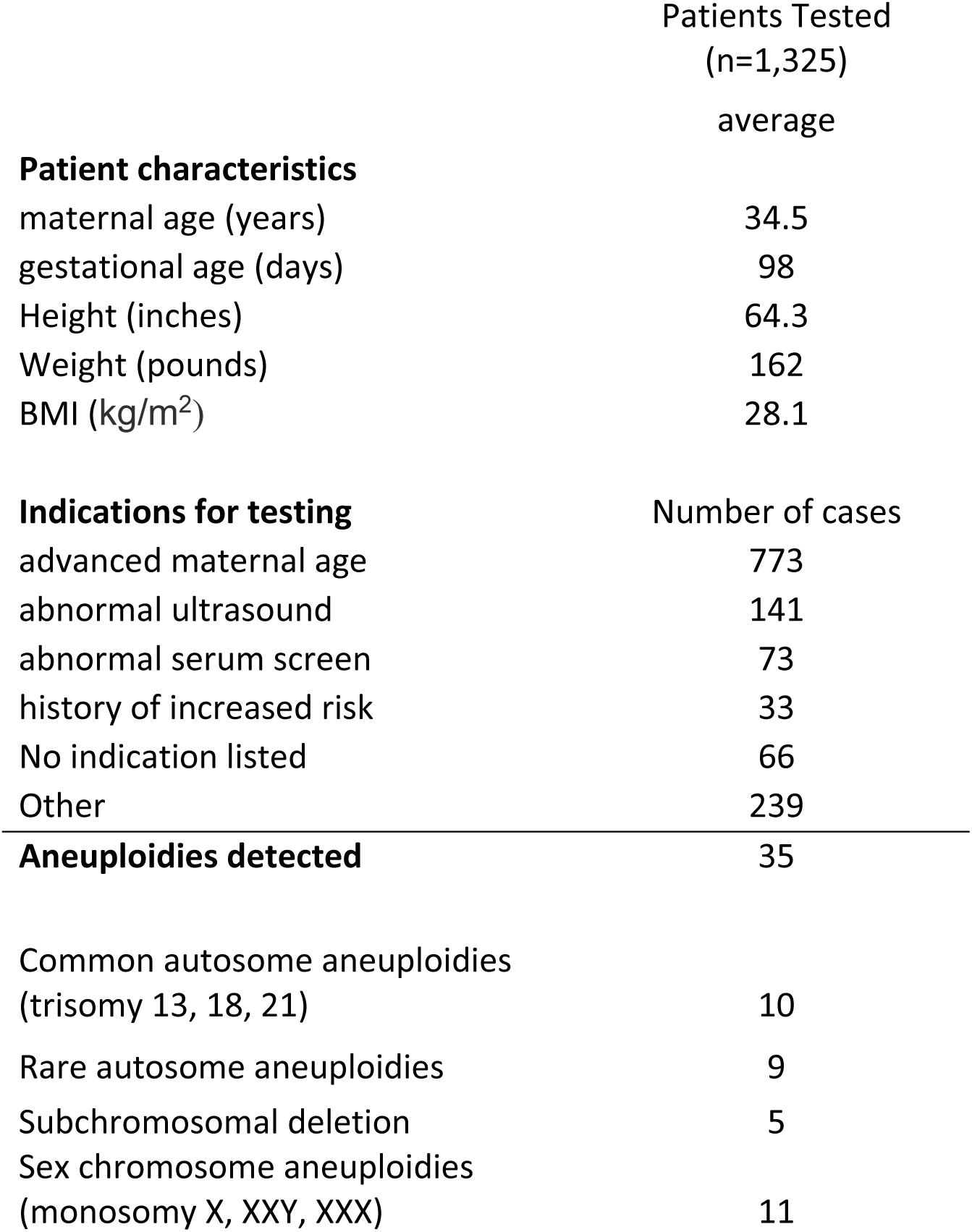
Patient demographics for the clinical implementation phase of our cfDNA prenatal screen. Data was collected at the same time as the specimen. A total of 1,481 samples were tested for 1,325 patients. These data do not include patient demographics for samples tested during the validation phase.

**Supplemental Table 2.** Quantities for CMV copies per mL plasma for positive samples identified by cfDNA and analyzed by qPCR. Specimens marked with (*) were identified as intermediate positive samples based on copy number in cfDNA sequencing, while specimen 34P02 (^#^) had fewer than 5µL of DNA remaining.

| **Sample Name** | **CMV**<br>**copies / ml** | **Beta-globin**<br>**copies / mL** |
| --- | --- | --- |
| **10P13** | 269.3 | 2096.6 |
| **111P08** | 11.6 | 1024.3 |
| **121R04** | 358.6 | 1324.3 |
| **34P02<sup>#</sup>** | 0.0 | 699.6 |
| **3P13** | 433.8 | 4084.9 |
| **80P04** | 108.4 | 8564.4 |
| **84P05** | 41.0 | 3840.8 |
| **92P02** | 26.2 | 1422.4 |
| **92R11** | 5.6 | 1878.9 |
| **104R** | 29.3 | 495.9 |
| **44P05\*** | 2.5 | 1017.5 |
| **96R12\*** | 0.4 | 1822.9 |

**Supplemental Table 3.**
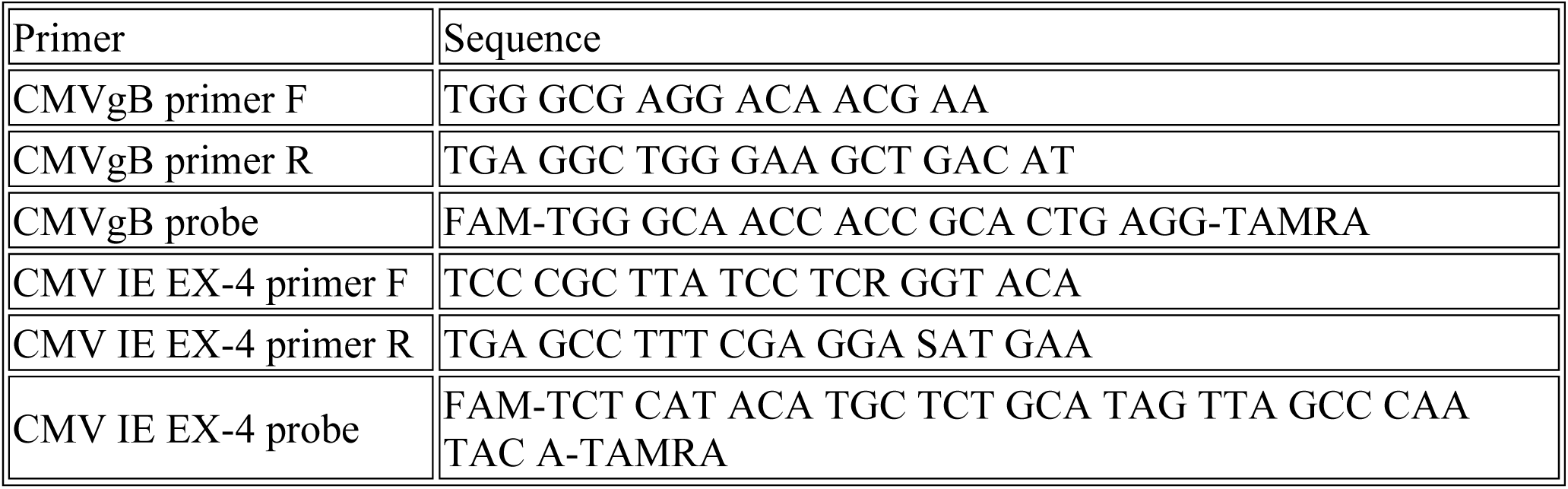
CMV qPCR primer and probe sequences.

## Supplemental Figures

**Supplemental Figure 1.**
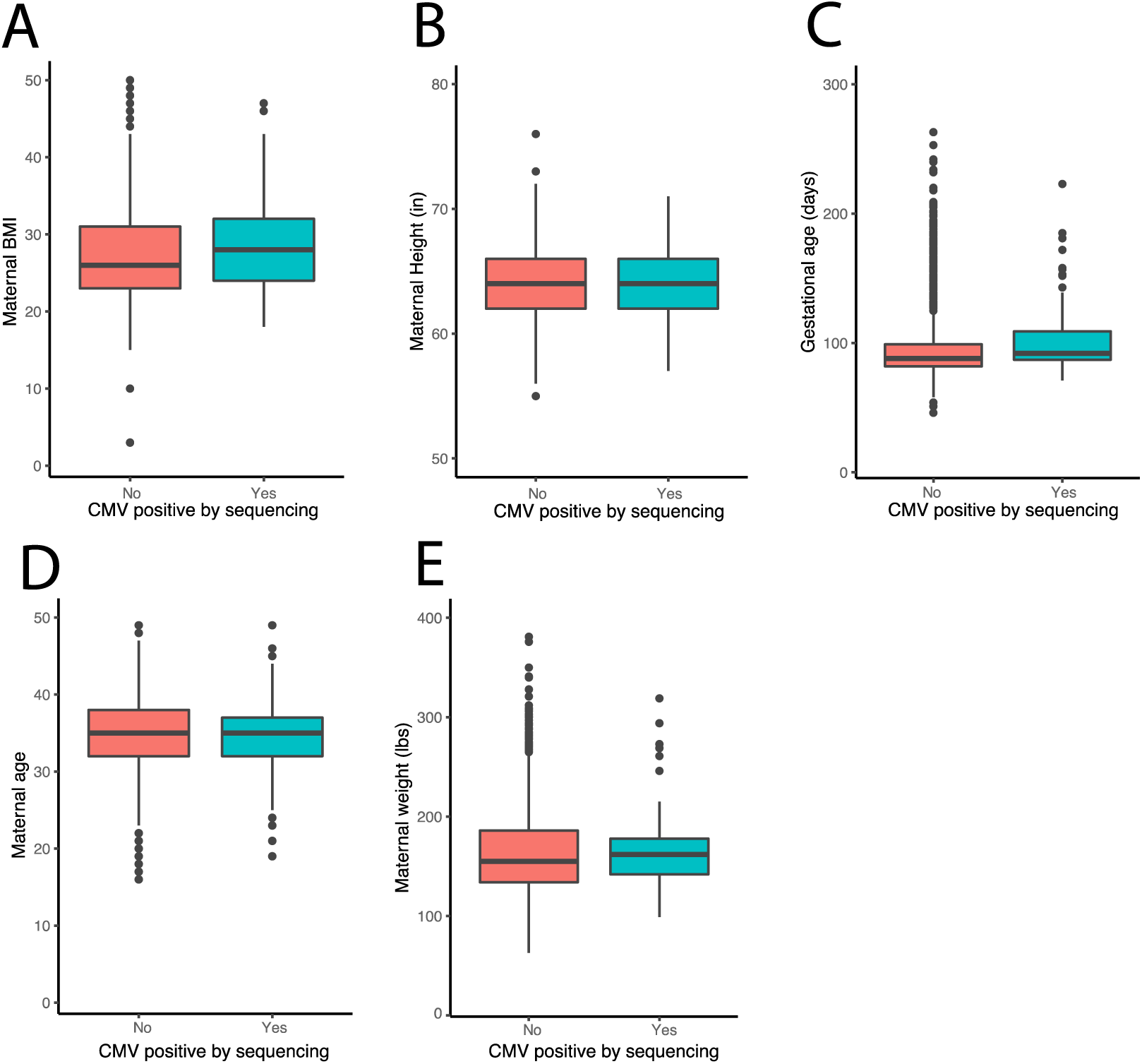
Comparison of maternal demographics based on CMV cfDNA positivity: A. Maternal BMI (kg/m^2^) B. Maternal Height (inches) C. Gestational age (days) D. Maternal age (years) E. Maternal Weight (pounds).

**Supplemental Figure 2.**
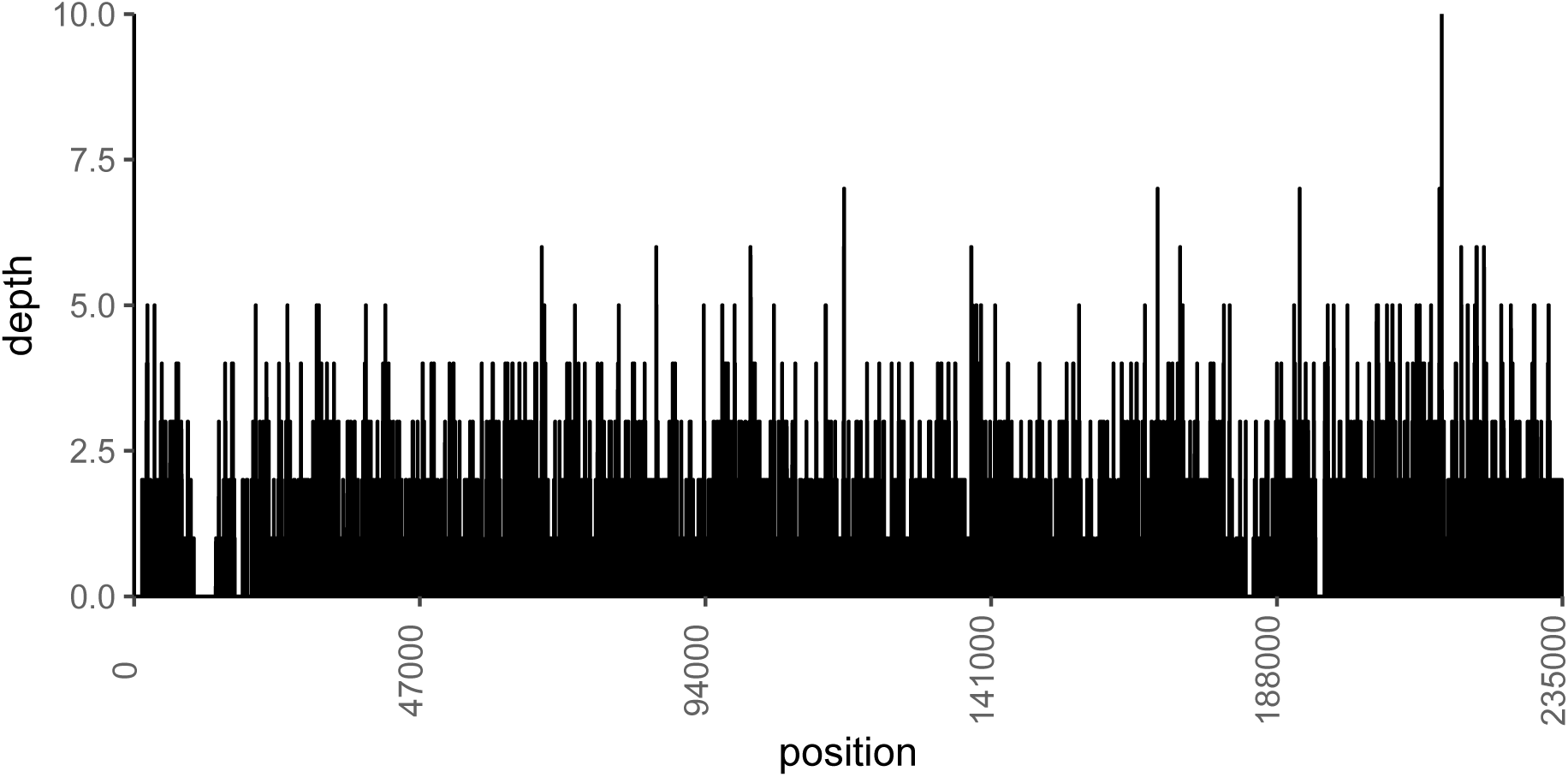
Read coverage of the CMV genome from sample 121R04. Reads were mapped to the human herpesvirus 5 Merlin strain reference genome.

## References

1. Kenneson A, Cannon MJ. 2007. Review and meta-analysis of the epidemiology of congenital cytomegalovirus (CMV) infection. Rev Med Virol 17:253–276.

2. Dollard SC, Grosse SD, Ross DS. 2007. New estimates of the prevalence of neurological and sensory sequelae and mortality associated with congenital cytomegalovirus infection. Rev Med Virol 17:355–363.

3. Society for Maternal-Fetal Medicine (SMFM), Hughes BL, Gyamfi-Bannerman C. 2016. Diagnosis and antenatal management of congenital cytomegalovirus infection. Am J Obstet Gynecol 214:B5–B11.

4. Ross SA, Boppana SB. 2005. Congenital cytomegalovirus infection: outcome and diagnosis. Semin Pediatr Infect Dis 16:44–49.

5. Adler SP. 2011. Screening for Cytomegalovirus during Pregnancy. Infect Dis Obstet Gynecol 2011.

6. Nigro G, Adler SP, La Torre R, Best AM, Congenital Cytomegalovirus Collaborating Group. 2005. Passive immunization during pregnancy for congenital cytomegalovirus infection. N Engl J Med 353:1350–1362.

7. Lazzarotto T, Guerra B, Lanari M, Gabrielli L, Landini MP. 2008. New advances in the diagnosis of congenital cytomegalovirus infection. J Clin Virol 41:192–197.

8. Adler SP. 2012. Editorial commentary: Primary maternal cytomegalovirus infection during pregnancy: do we have a treatment option? Clin Infect Dis 55:504–506.

9. Lazzarotto T, Guerra B, Gabrielli L, Lanari M, Landini MP. 2011. Update on the prevention, diagnosis and management of cytomegalovirus infection during pregnancy. Clin Microbiol Infect 17:1285–1293.

10. Enders G, Bäder U, Lindemann L, Schalasta G, Daiminger A. 2001. Prenatal diagnosis of congenital cytomegalovirus infection in 189 pregnancies with known outcome. Prenat Diagn 21:362–377.

11. Donner C, Liesnard C, Brancart F, Rodesch F. 1994. Accuracy of amniotic fluid testing before 21 weeks’ gestation in prenatal diagnosis of congenital cytomegalovirus infection. Prenat Diagn 14:1055–1059.

12. Liesnard C, Donner C, Brancart F, Gosselin F, Delforge ML, Rodesch F. 2000. Prenatal diagnosis of congenital cytomegalovirus infection: prospective study of 237 pregnancies at risk. Obstet Gynecol 95:881–888.

13. Boppana SB, Ross SA, Novak Z, Shimamura M, Tolan RW, Palmer AL, Ahmed A, Michaels MG, Sánchez PJ, Bernstein DI, Britt WJ, Fowler KB, Study for the NI on D and OCDC and HMS (CHIMES). 2010. Dried Blood Spot Real-time Polymerase Chain Reaction Assays to Screen Newborns for Congenital Cytomegalovirus Infection. JAMA 303:1375–1382.

14. Kimberlin DW, Lin C-Y, Sánchez PJ, Demmler GJ, Dankner W, Shelton M, Jacobs RF, Vaudry W, Pass RF, Kiell JM, Soong S, Whitley RJ, National Institute of Allergy and Infectious Diseases Collaborative Antiviral Study Group. 2003. Effect of ganciclovir therapy on hearing in symptomatic congenital cytomegalovirus disease involving the central nervous system: a randomized, controlled trial. J Pediatr 143:16–25.

15. Fowler KB, McCollister FP, Dahle AJ, Boppana S, Britt WJ, Pass RF. 1997. Progressive and fluctuating sensorineural hearing loss in children with asymptomatic congenital cytomegalovirus infection. J Pediatr 130:624–630.

16. Misono S, Sie KCY, Weiss NS, Huang M, Boeckh M, Norton SJ, Yueh B. 2011. Congenital Cytomegalovirus Infection in Pediatric Hearing Loss. Arch Otolaryngol Head Neck Surg 137:47–53.

17. Lo YM, Corbetta N, Chamberlain PF, Rai V, Sargent IL, Redman CW, Wainscoat JS. 1997. Presence of fetal DNA in maternal plasma and serum. Lancet 350:485–487.

18. Lo YM, Tein MS, Lau TK, Haines CJ, Leung TN, Poon PM, Wainscoat JS, Johnson PJ, Chang AM, Hjelm NM. 1998. Quantitative analysis of fetal DNA in maternal plasma and serum: implications for noninvasive prenatal diagnosis. Am J Hum Genet 62:768–775.

19. Chesnais V, Ott A, Chaplais E, Gabillard S, Pallares D, Vauloup-Fellous C, Benachi A, Costa J-M, Ginoux E. 2018. Using massively parallel shotgun sequencing of maternal plasmatic cell-free DNA for cytomegalovirus DNA detection during pregnancy: a proof of concept study. Sci Rep 8:4321.

20. Wickham H. 2009. Ggplot2: Elegant Graphics for Data Analysis, 2nd ed. Springer Publishing Company, Incorporated.

21. Sedlak RH, Hill JA, Nguyen T, Cho M, Levin G, Cook L, Huang M-L, Flamand L, Zerr DM, Boeckh M, Jerome KR. 2016. Detection of Human Herpesvirus 6B (HHV-6B) Reactivation in Hematopoietic Cell Transplant Recipients with Inherited Chromosomally Integrated HHV-6A by Droplet Digital PCR. J Clin Microbiol 54:1223–1227.

22. Society for Maternal-Fetal Medicine (SMFM), Hughes BL, Gyamfi-Bannerman C. 2016. Diagnosis and antenatal management of congenital cytomegalovirus infection. Am J Obstet Gynecol 214:B5–B11.

23. American College of Obstetricians and Gynecologists. 2015. Practice bulletin no. 151: Cytomegalovirus, parvovirus B19, varicella zoster, and toxoplasmosis in pregnancy. Obstet Gynecol 125:1510–1525.

24. Hayden RT, Preiksaitis J, Tong Y, Pang X, Sun Y, Tang L, Cook L, Pounds S, Fryer J, Caliendo AM. 2015. Commutability of the First World Health Organization International Standard for Human Cytomegalovirus. J Clin Microbiol 53:3325–3333.

25. Preiksaitis JK, Hayden RT, Tong Y, Pang XL, Fryer JF, Heath AB, Cook L, Petrich AK, Yu B, Caliendo AM. 2016. Are We There Yet? Impact of the First International Standard for Cytomegalovirus DNA on the Harmonization of Results Reported on Plasma Samples. Clin Infect Dis 63:583–589.

26. Boom R, Sol CJA, Schuurman T, van Breda A, Weel JFL, Beld M, ten Berge IJM, Wertheim-van Dillen PME, de Jong MD. 2002. Human Cytomegalovirus DNA in Plasma and Serum Specimens of Renal Transplant Recipients Is Highly Fragmented. J Clin Microbiol 40:4105–4113.

27. Tong Y, Pang XL, Mabilangan C, Preiksaitis JK. 2017. Determination of the Biological Form of Human Cytomegalovirus DNA in the Plasma of Solid-Organ Transplant Recipients. J Infect Dis 215:1094–1101.

28. Liu S, Huang S, Chen F, Zhao L, Yuan Y, Francis SS, Fang L, Li Z, Lin L, Liu R, Zhang Y, Xu H, Li S, Zhou Y, Davies RW, Liu Q, Walters RG, Lin K, Ju J, Korneliussen T, Yang MA, Fu Q, Wang J, Zhou L, Krogh A, Zhang H, Wang W, Chen Z, Cai Z, Yin Y, Yang H, Mao M, Shendure J, Wang J, Albrechtsen A, Jin X, Nielsen R, Xu X. 2018. Genomic Analyses from Non-invasive Prenatal Testing Reveal Genetic Associations, Patterns of Viral Infections, and Chinese Population History. Cell 175:347-359.e14.

